# gPlace: fast, memory-bounded phylogenetic placement with results identical to pplacer

**DOI:** 10.64898/2026.11.23.768714

**Authors:** Pierre-Alain Chaumeil, Donovan H. Parks

## Abstract

Phylogenetic placement methods such as pplacer attach query sequences to a fixed reference tree, thereby establishing their evolutionary relationships to the reference sequences. These relationships can then be used to infer taxonomic classifications or likely metabolic functions. gPlace is a drop-in replacement for pplacer that produces identical results while reducing memory requirements by >3-fold and computation time by >30-fold under typical usage. These improvements are achieved by storing less data per node and streamlining the calculations used to position each query sequence. gPlace can also run within a user-specified memory limit by processing only a portion of the reference tree at a time. Consequently, it can efficiently place sequences on large reference trees that would be impractical for pplacer to process. We demonstrate the importance of these improvements by evaluating gPlace as a replacement for pplacer in GTDB-Tk and CheckM, two tools widely used in genome-resolved metagenomic studies. gPlace is released under the GNU General Public License v3.0 and is available at https://github.com/cmc-aau/gPlace.

## Statement of Need

Phylogenetic placement methods such as pplacer (Matsen et al., 2010) and the Evolutionary Placement Algorithm (EPA; Berger et al., 2011) were developed to place short sequences into single-gene reference trees (e.g., 16S rRNA) and have since been used to place genome assemblies into trees inferred from concatenated marker genes (Parks et al., 2015). Longer alignments, combined with rapidly growing reference trees, have made placement increasingly demanding in memory and runtime. GTDB-Tk (Chaumeil et al., 2020), a widely used bioinformatics tool for determining the taxonomic classification of bacterial and archaeal genome assemblies, illustrates this challenge. For bacteria, it classifies genomes by using pplacer to place them into a Genome Taxonomy Database (GTDB; Parks et al., 2026) reference tree inferred from a ∼5,000-column concatenated alignment of 120 marker genes. This tree has grown by ∼34% annually to 189,801 species in R11-RS232 (GTDB release 11, RefSeq release 232), and placing genomes into a tree of this size with pplacer requires 912 GiB of memory. GTDB-Tk v2 (Chaumeil et al., 2022) reduces this memory requirement by placing bacterial genomes first into a backbone tree and then into class-level subtrees. However, the largest class-level tree now requires 140 GiB, bringing GTDB-Tk’s peak memory requirement to ∼190 GiB when run with 64 processes. This is beyond the resources readily available to many researchers. gPlace addresses this challenge with a computationally efficient placement algorithm that runs within a user-specified memory bound while producing placements identical to those of pplacer.

## State of the Field

Phylogenetic placement was popularized by two maximum-likelihood (ML) tools, pplacer and EPA, and later work has largely aimed to reduce their computational cost (**Figure 1A**). EPA-ng (Barbera et al., 2019) is a parallel reimplementation of EPA that reduces runtime but requires more memory than pplacer (**Figure 1B**). GTDB-Tk v2 (Chaumeil et al., 2022), SCAMPP (Wedell et al., 2023), BSCAMPP (Wedell et al., 2025), and Splicer (Markin & Anderson, 2026) reduce memory and, in most cases, runtime by applying pplacer or EPA-ng to subtrees rather than the full reference tree; however, placements can differ from full-tree placement (**Figure 1C**). Other tools replace ML with faster criteria. APPLES-2 (Balaban et al., 2022) places queries by distance, and UShER (Turakhia et al., 2021) by parsimony. APPLES-2 scales to large trees but gives different placements from ML methods (**Figure 1C**), while UShER requires mutation-annotated nucleotide trees, making it less flexible than other placement methods. In contrast, gPlace reduces computational requirements while providing results identical to those of pplacer, allowing it to serve as a no-compromise, drop-in replacement. For example, running gPlace with a memory bound of 64 GiB resulted in a 35.7-fold decrease in computation time and a 15.9-fold decrease in memory usage relative to pplacer (**Figure 1B**).

**Figure 1.**
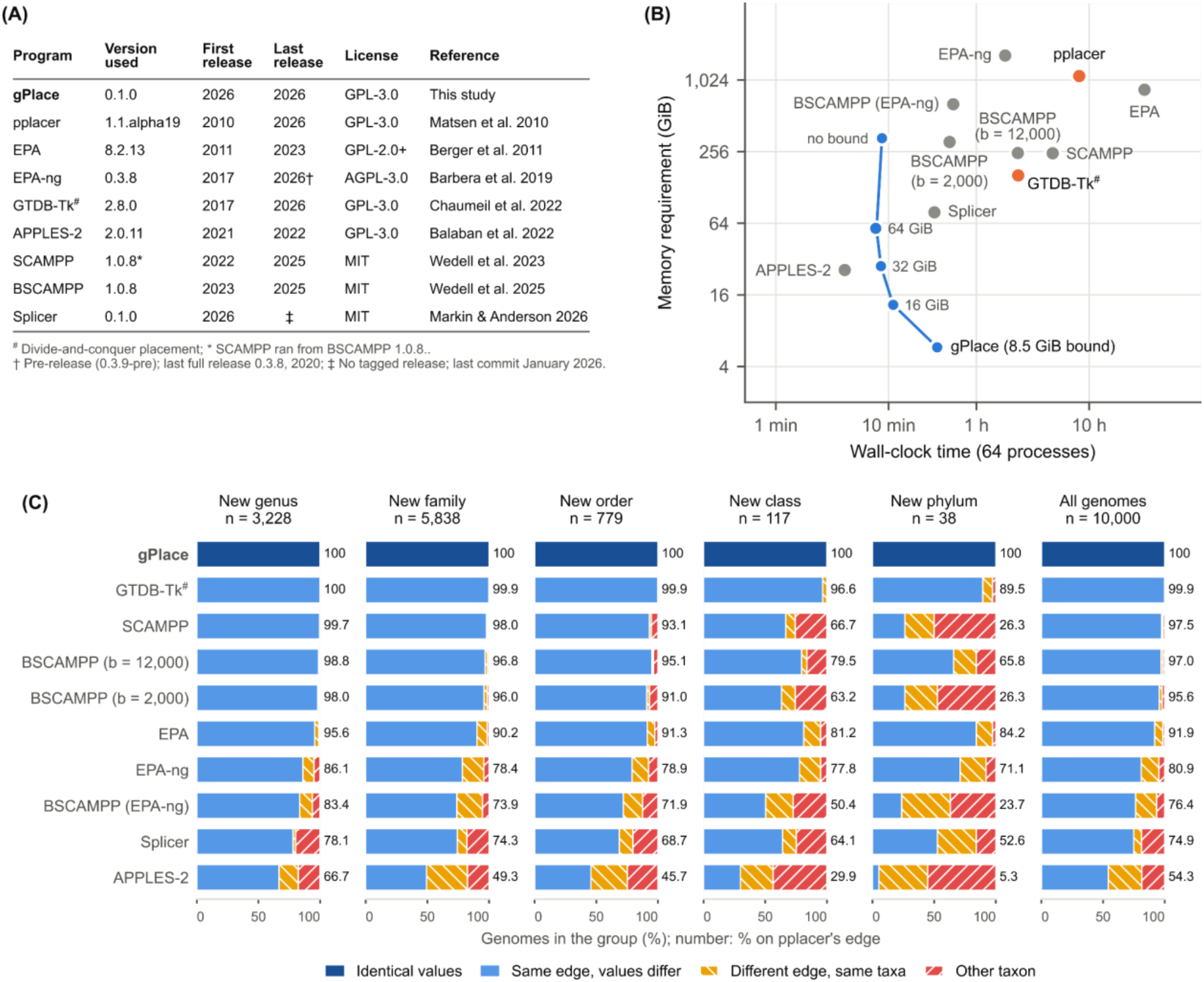
Comparison of phylogenetic placement tools. Each tool was used to place 10,000 GlobDB release 232 (Speth et al., 2025) bacterial genomes not assigned to a GTDB R11-RS232 species into the GTDB R11-RS232 bacterial reference tree (189,801 leaves; 5,010-column alignment). All tools were run with 64 processes on an AMD EPYC 9565 server with 2 TiB of RAM. (**A**) Placement tools evaluated, with versions, release dates, and licenses. (**B**) Wall-clock time and peak memory requirement of each tool. gPlace was run with maximum memory bounds of 8.5, 16, 32, and 64 GiB; BSCAMPP was run using pplacer on subtrees of 2,000 (default) and 12,000 leaves and using EPA-ng with default settings; UShER was not evaluated as it requires mutation-annotated nucleotide trees. (**C**) Placements produced by each tool relative to those obtained with pplacer. Results are stratified by taxonomic novelty and indicate whether a placement is identical (i.e., same edge and likelihood), on the same edge but with a different position and/or likelihood, on a different edge within the same taxon, or on an edge within a different taxon. The number to the right of each bar is the percentage of genomes placed on the same edge as pplacer (i.e., with or without identical likelihood).

## Software Design

### Improvements to Reduce Memory Requirements

The memory required by pplacer is dominated by the partial likelihood vectors it caches for every node of the reference tree (**Figure 2A**). Each node holds three vectors: i) a distal vector, the partial likelihood of the subtree below the node; ii) a proximal vector, the partial likelihood of the rest of the tree; and iii) a supernode vector, the distal and proximal vectors combined at the midpoint of the edge above the node, which pplacer uses to rank candidate edges before optimizing edge lengths. For a reference tree with *N* internal nodes, *L* leaf nodes, and an alignment of length *A*, pplacer requires 3 × (*N* + *L*) × *A* × 20 × 8 bytes, as each vector requires 8 bytes for each of the 20 amino-acid states at each alignment column.

**Figure 2.**
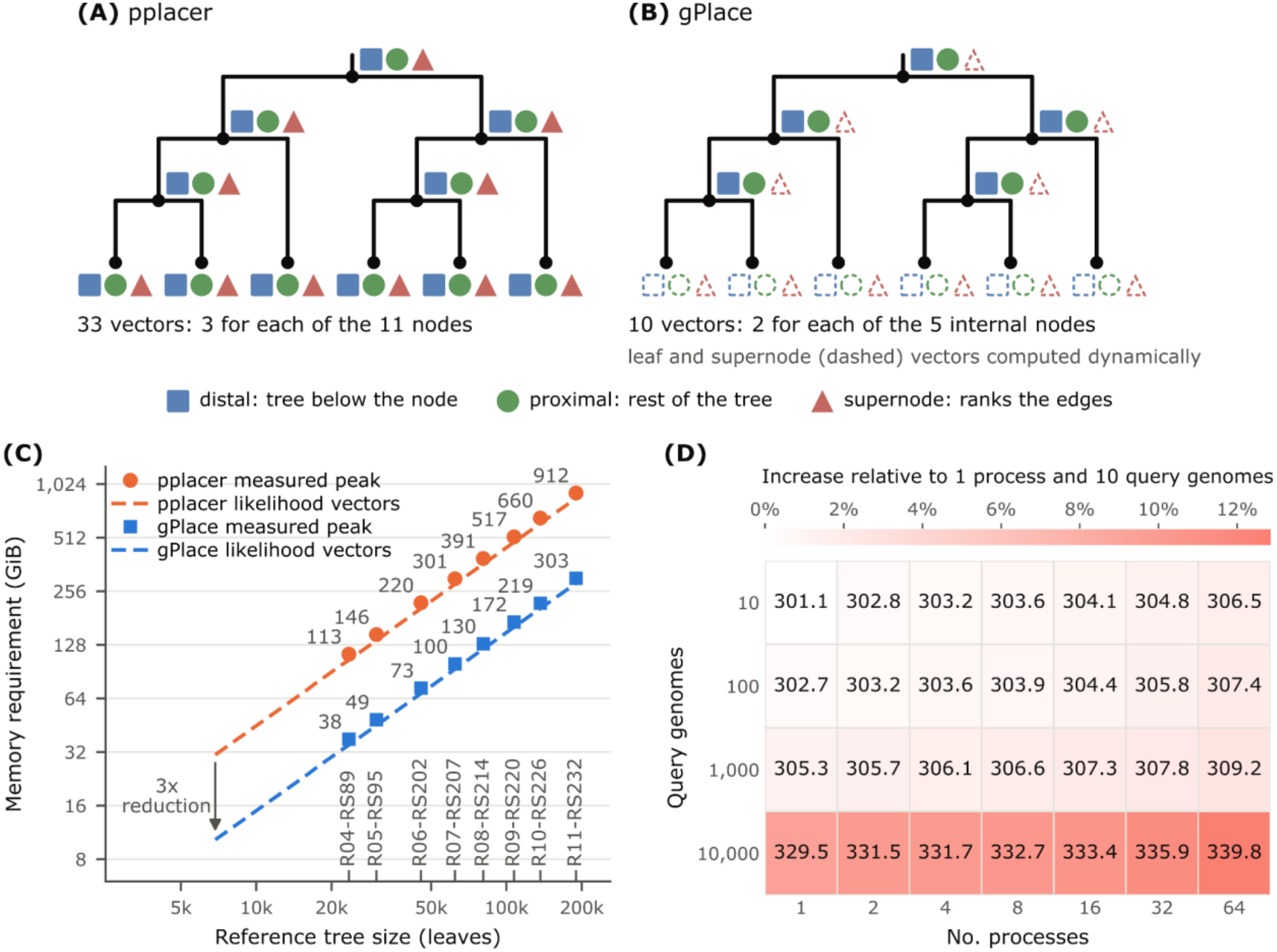
Likelihood-vector storage in pplacer and gPlace. (**A**) pplacer caches distal, proximal, and supernode vectors for every node. (**B**) gPlace caches distal and proximal vectors for internal nodes only; leaf and supernode vectors (dashed) are rebuilt when needed. (**C**) Calculated memory of the cached vectors (dashed lines) and measured peak memory (points) against the number of leaves in GTDB bacterial reference trees, all of which have ∼5,000-column alignments. Testing was performed for 100 query genomes with a single process on a 1 TiB AMD EPYC 7402 server. (**D**) Peak memory (GiB) of gPlace on the GTDB R11-RS232 bacterial tree as a function of the number of processes and query genomes.

gPlace keeps only the distal and proximal vectors, and only for internal nodes (**Figure 2B**). This reduces memory requirements to 2 × *N* × *A* × 20 × 8 bytes, a 3-fold reduction on bifurcating trees (**Figure 2C**), and more on multifurcating trees. The vectors that are no longer stored are cheap to rebuild. A leaf’s distal vector is its aligned sequence, a leaf’s proximal vector is its parent’s proximal vector multiplied by its siblings’ distal vectors, and supernode vectors are computed eight edges at a time as needed, then discarded. These vectors are rebuilt using the same operations in the same order as in pplacer, so they are identical to those pplacer stores. Although pplacer and gPlace require additional memory for a per-column scaling exponent (*A* × 8 bytes per vector), the alignment, the reference tree, query genomes, and per-process overhead, memory requirements are dominated by the likelihood vectors (**Figures 2C** and **2D**).

### Placement with a Maximum Memory Bound

The distal and proximal likelihood vectors can still exceed available memory for large reference trees. To address this, gPlace can partition the reference tree into connected pieces, termed tiles, and hold only one tile in memory at a time, with tile size chosen so that placement stays within a user-specified memory limit (**Figure 3**). A tile meets the rest of the tree only at the edge above its root and at the roots of the tiles below it, so its vectors can be rebuilt exactly from two boundary vectors per tile root: the distal vector of the subtree below it, saved during a bottom-up pass over the tiles, and the proximal vector of the rest of the tree, saved during a top-down pass. Every likelihood computed within a tile is therefore still a full-tree likelihood, unlike in divide-and-conquer methods, which place queries on subtrees (Chaumeil et al., 2022; Wedell et al., 2023, 2025; Markin & Anderson, 2026).

**Figure 3.**
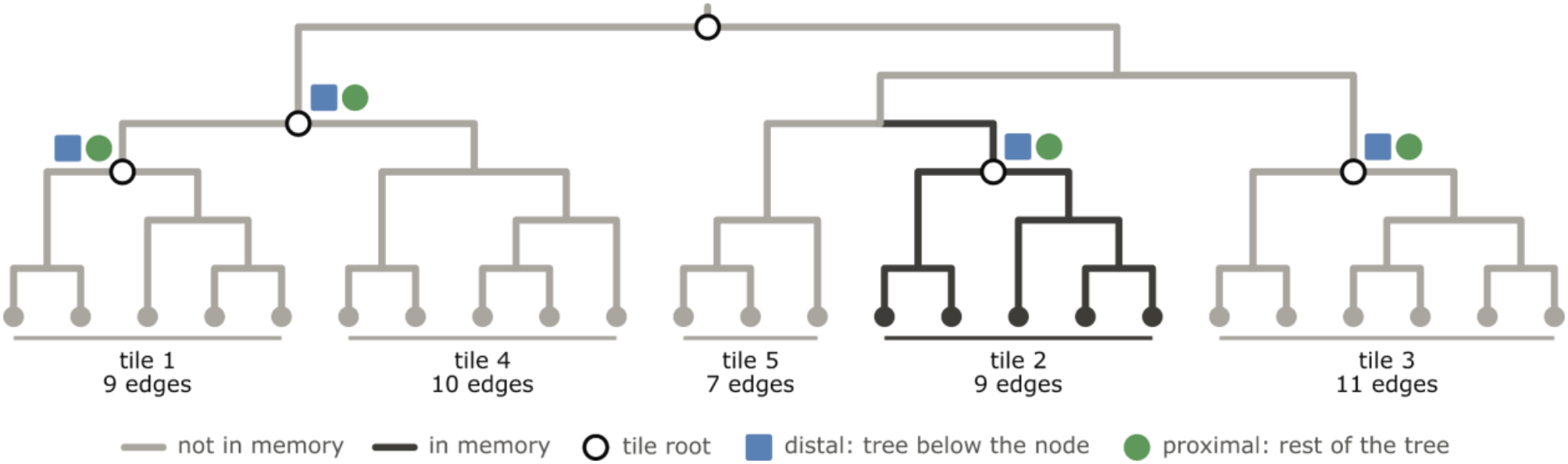
Tiling the reference tree allows phylogenetic placement within a user-specified memory bound. A 24-leaf reference tree is partitioned into five connected tiles, each with at most 12 edges, numbered in the order the bottom-up pass builds them. Two boundary vectors are stored for every tile root except the tree root: a distal vector for the subtree below it and a proximal vector for the rest of the tree. These are enough to rebuild any tile’s likelihood vectors exactly, so only one tile (tile 2 in this example, shown in black) needs to be in memory at a time.

For each query sequence, pplacer ranks all edges by a fast approximate score and then works down this ranking to optimize the query’s position on each edge and the length of the new edge connecting it to the tree. This continues until a set number of edges have proven clearly worse than the best edge found so far. Finally, the best few placements are further refined. Applied to a tiled tree, this per-query search would move constantly between tiles. gPlace instead processes all queries one tile at a time, ranking edges by scoring every query against the edges of each tile and keeping each query’s top-ranked edges. Because the optimization on each candidate edge always starts from the same initial values, its result does not depend on the order in which edges are evaluated. The search for the best edges can therefore proceed in rounds: each round evaluates the next few ranked edges of every query still searching, tile by tile, then applies the same stopping rule to the accumulated results, giving the same placements as pplacer’s sequential search. The best placements are then refined tile by tile. Tiling trades additional computation for reduced memory, as the likelihood vectors of each tile must be recomputed once to rank edges, once per search round, and once to refine placements (**Figure 1B**).

### Improvements to Reduce Computation Time

Scoring and ranking the edges of the reference tree, the first step of pplacer’s search, dominates its runtime. For example, when placing 10,000 genomes on the GTDB R08-RS214 bacterial tree (161,575 edges) with 64 processes, scoring and ranking took 81% (3 h 40 min) of pplacer’s total runtime (4 h 30 min; **Figure 4D**). gPlace reduces computation time primarily by optimizing this step, placing the same genomes 51-fold faster (5 min 20 s).

**Figure 4.**
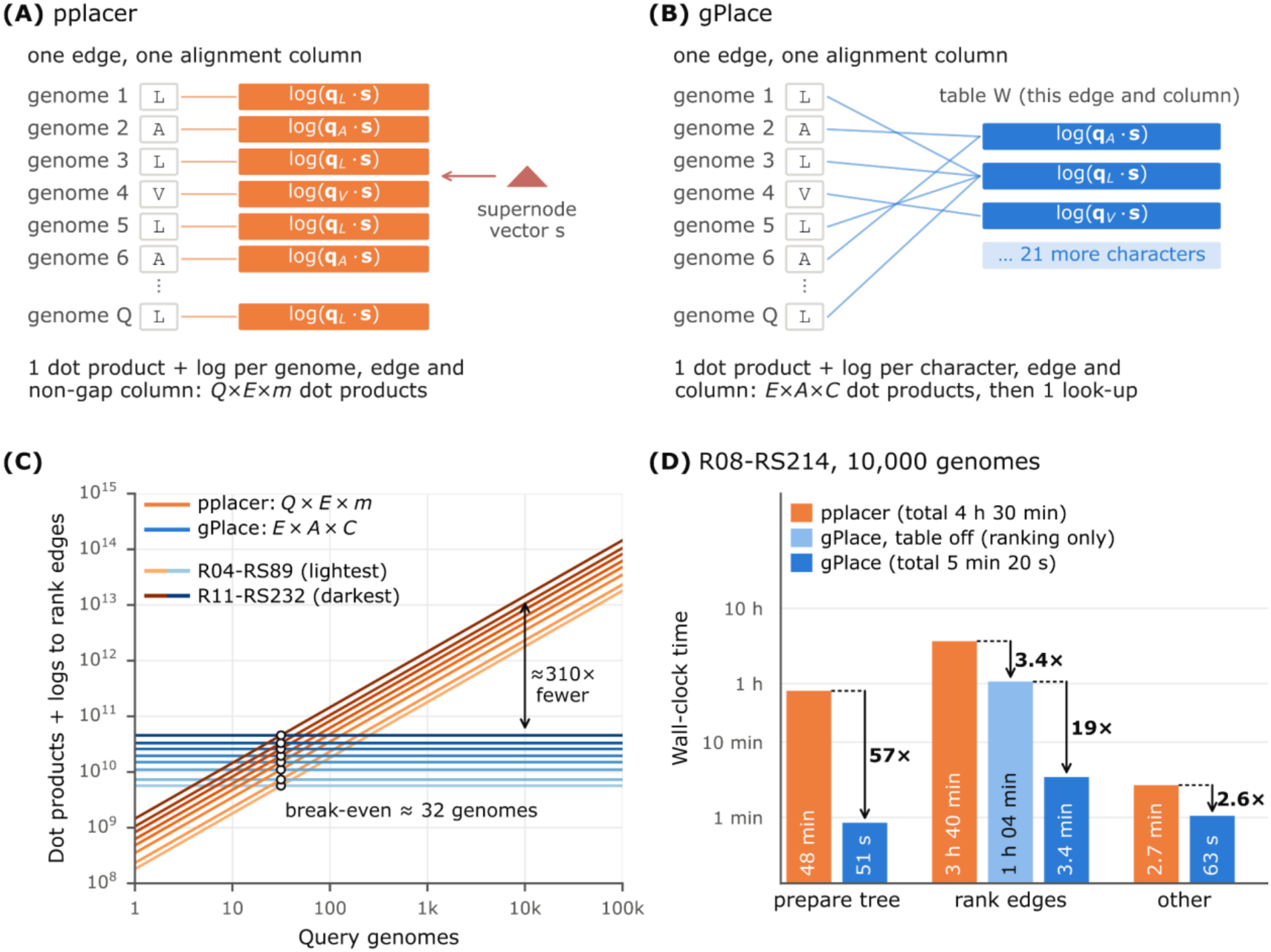
gPlace substantially reduces computation time by changing how edges are scored and ranked. (**A**) pplacer ranks the edges for one genome at a time, computing log(*q* · *s*) from the genome’s amino acid (as vector *q*) and the edge’s supernode vector (*s*) for every genome, edge, and non-gap column. (**B**) gPlace ranks the edges for all genomes together, computing this value once per possible character (table *W*) for each edge and column so that each genome sums one table entry per column. (**C**) Dot products and logarithms needed to rank all edges of the GTDB R04-RS89 (lightest lines; 46,913 edges) to R11-RS232 (darkest; 379,593 edges) bacterial trees, assuming 75% of columns are non-gap, as in R11-RS232. Circles mark the break-even points, and the arrow shows the reduction at 10,000 genomes for R11-RS232. (**D**) Wall-clock time per stage for placing 10,000 GlobDB bacterial genomes not assigned to a GTDB R11-RS232 species into the GTDB R08-RS214 tree using 64 processes on an AMD EPYC 7402 server with 1 TiB of RAM. The pplacer and gPlace bars each sum to that tool’s total time, and “table off” is gPlace without the per-edge table optimization.

The score used to rank a genome on an edge approximates the log-likelihood of attaching it there. It is a sum over alignment columns of log(*q* · *s*), where *s* is the edge’s supernode vector and *q* is the equivalent vector for the genome, derived from its amino acid in that column and the substitution probabilities along a new edge of fixed initial length. pplacer ranks the edges for one genome at a time, computing this 20-term dot product and logarithm for every edge and non-gap column (**Figure 4A**), and so passes over the supernode vectors of the whole tree once per genome. gPlace instead builds the supernode vectors of a few edges at a time and scores all genomes against them before moving on (**Figure 4B**).

Scoring all genomes together also allows per-character contributions to be tabulated, as in the preplacement lookup table described for EPA-ng (Barbera & Stamatakis, 2021). Whereas EPA-ng stores this table for every edge, gPlace builds it for a few edges at a time and then discards it, so it adds no appreciable memory. Within a given alignment column, *q* depends only on the genome’s amino acid, so all genomes with the same amino acid in that column receive the same contribution. gPlace therefore computes, for each edge, a table holding the contribution of each of the *C* = 24 possible characters (20 amino acids and the ambiguity codes B, J, Z, and X) at each column and scores each genome by summing one table entry per column (**Figure 4B**). For *Q* genomes with a mean of *m* non-gap columns, a tree with *E* edges, and an alignment of *A* columns, pplacer computes *Q* × *E* × *m* dot products and logarithms, whereas gPlace computes *E* × *A* × *C* regardless of *Q* (**Figure 4C**). The table requires fewer operations once *Q* exceeds *A* × *C* / *m* (e.g., ∼32 genomes for GTDB alignments, where genomes cover ∼75% of columns) and *Q* × *m* / (*A* × *C*)-fold fewer above this (e.g., ∼310-fold for 10,000 genomes; **Figure 4C**).

Without the table optimization, gPlace required 1 h 4 min to score and rank edges, 3.4-fold faster than pplacer, and the table made ranking a further 19-fold faster (3 min 24 s; 64-fold overall; **Figure 4D**). Most of the remaining difference lies in preparing the reference tree (48 min for pplacer, 51 s for gPlace), as gPlace computes only the distal and proximal vectors of internal nodes. Faster edge-length optimization and other changes account for less than 1% of the time saved.

## Research Impact Statement

We demonstrate the benefits of gPlace as a drop-in replacement for pplacer by evaluating its impact on two widely used bioinformatics tools, GTDB-Tk (Chaumeil et al., 2020, 2022) and CheckM (Parks et al., 2015).

GTDB-Tk has >8,900 citations (Google Scholar, 18 Nov. 2026) and is integrated into bioinformatics pipelines (e.g., ATLAS; Kieser et al., 2020), resources (e.g., MGnify; Gurbich et al., 2023), and computing platforms (e.g., KBase; Chivian et al., 2023). Placing query genomes into the GTDB bacterial reference tree with pplacer is the main runtime and memory bottleneck of GTDB-Tk, which the divide-and-conquer strategy of GTDB-Tk v2 only partially mitigates (**Figure 1B**). To assess the impact of gPlace, we ran GTDB-Tk v2.8.0 with the GTDB R11-RS232 bacterial tree on 10,000 query genomes using four placement configurations (**Figure 5A**): pplacer on the full tree, pplacer with divide-and-conquer, and gPlace on the full tree with and without a specified memory bound. All runs used 64 processes on an AMD EPYC 7402 server with 1 TiB of RAM. GTDB-Tk with full-tree pplacer placement required 19 h 4 min, 241.1 GiB of memory, and an 893 GiB memory-mapped scratch file. Without the scratch file, pplacer has insufficient memory to run with 64 processes and falls back to a single process requiring 912 GiB. As few researchers have access to machines with ∼1 TiB of memory, the scratch-file results reflect how most users would run GTDB-Tk when requiring full-tree pplacer results. GTDB-Tk with divide-and-conquer placement required 6 h 6 min and 187.3 GiB, with 2 of 10,000 genomes receiving a different classification than with full-tree placement. In contrast, GTDB-Tk with gPlace produced placements identical to full-tree pplacer in 1 h 6 min using 333.7 GiB and no scratch file. Limiting gPlace to 32 GiB increased the runtime to 1 h 16 min and reduced GTDB-Tk’s peak memory to 39.3 GiB, nearly 5-fold less time and memory than divide-and-conquer placement. Notably, gPlace is no longer the bottleneck in GTDB-Tk. Other steps in the workflow dominate the wall-clock time and, once gPlace is capped below 40 GiB, the peak memory use as well. gPlace makes full-tree placement into the GTDB reference tree practical on readily available hardware and removes the need for the divide- and-conquer approach in GTDB-Tk.

**Figure 5.**
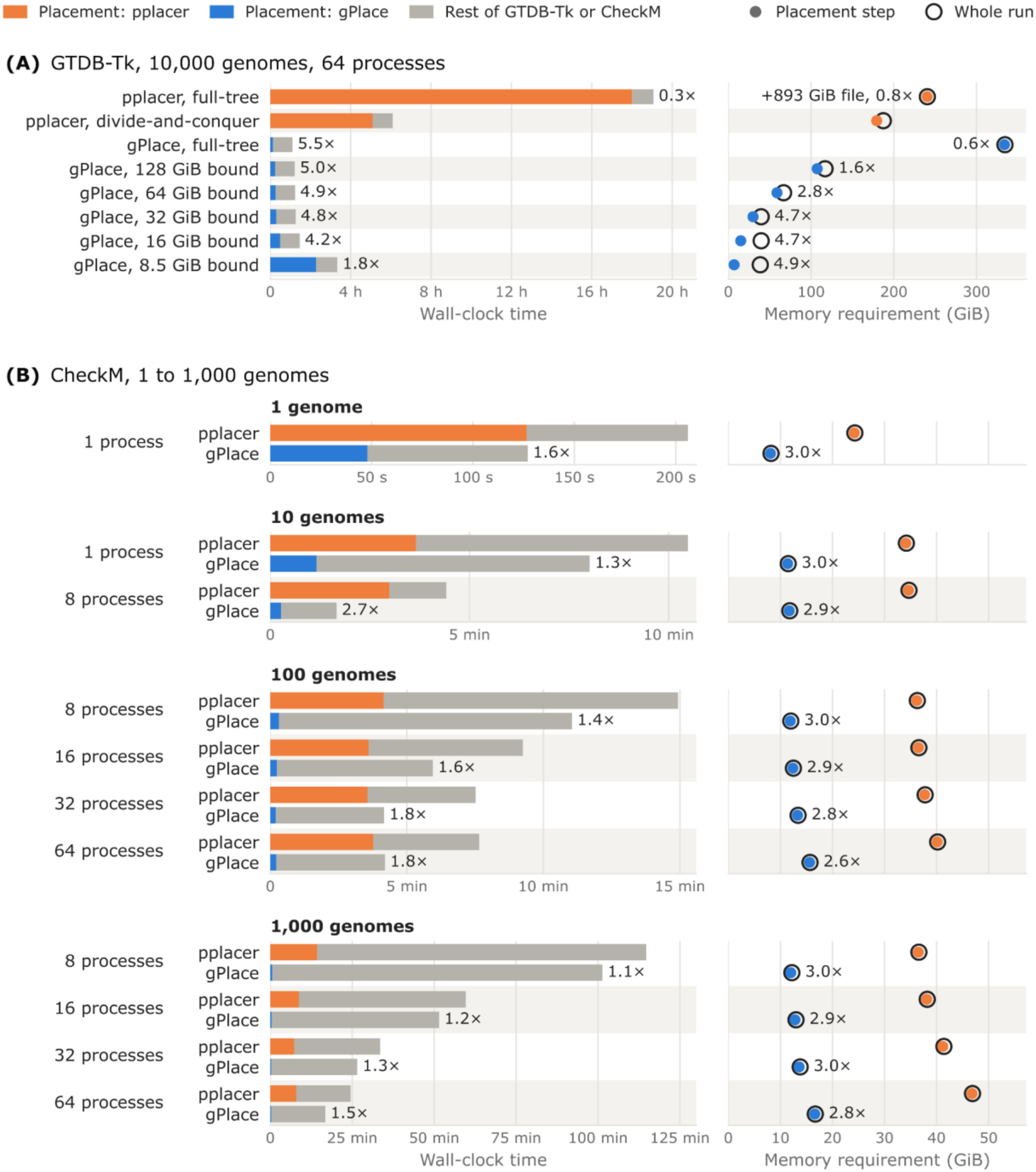
Run time and memory use of GTDB-Tk and CheckM using pplacer or gPlace for phylogenetic placement. Bars show wall-clock time, split into the placement step (colored) and the rest of the run (grey). Points show peak memory of the placement step (filled) and of the whole run (open). (**A**) GTDB-Tk classifying 10,000 bacterial genomes against the GTDB R11-RS232 reference tree using 64 processes. Query genomes were randomly selected from GlobDB release 232 genomes not assigned to an R11-RS232 species. pplacer was run on the full tree or with GTDB-Tk’s divide-and-conquer approach; gPlace was run on the full tree without a memory limit or with the indicated memory bound. Fold values (×) are relative to pplacer with divide-and-conquer. **(B)** CheckM applied to 1, 10, 100, or 1,000 genomes randomly selected from those in (A), with 1 to 64 processes. Fold values (×) are relative to the pplacer run with the same number of genomes and processes.

CheckM estimates the completeness and contamination of genome assemblies. It has >14,400 citations (Google Scholar, 18 Nov. 2026) and is integrated into numerous bioinformatics resource, including RefSeq at NCBI (Haft et al., 2024). CheckM relies on phylogenetic placement to establish a suitable set of marker genes for estimating the quality of a genome assembly. The reference tree used by CheckM is much smaller than those used by GTDB-Tk, consisting of only 5,242 leaves though with a longer alignment of 6,988 amino acids. Replacing pplacer with gPlace in CheckM v1.3.2 reduced wall-clock time 1.1-to 2.7-fold, depending on the number of query genomes and processes, and reduced peak memory approximately 3-fold in nearly all configurations (**Figure 5B**). Together, these results demonstrate that tools that rely on pplacer benefit from replacing pplacer with gPlace.

## AI Usage Disclosure

The gPlace codebase, unit tests, validation scripts, and documentation were written using Claude Opus 5.5. The authors specified the software design requirements for gPlace, proposed research directions, and decided which improvements to incorporate. This paper was drafted with Claude Opus 5.5 and heavily revised by the authors. The authors reviewed, edited, and validated all AI-assisted outputs.

## Acknowledgements

We thank the authors of pplacer, guppy, and taxtastic, from which gPlace is derived, and of GSL, CORE-MATH, NumPy, and tqdm, on which gPlace depends.

## Funding

This study was funded by the Novo Nordisk Foundation (NNF25SA0105111), the Australian Research Council (Discovery Project DP220100900), and The University of Queensland (strategic funding).

## Notes

### Competing Interest Statement

The authors have declared no competing interest.

